# scKAN: Interpretable Single-cell Analysis for Cell-type-specific Gene Discovery and Drug Repurposing via Kolmogorov-Arnold Networks

**DOI:** 10.1101/2025.02.04.636408

**Authors:** Haohuai He, Zhenchao Tang, Guanxing Chen, Fan Xu, Yao Hu, Yinglan Feng, Jibin Wu, Yu-An Huang, Zhi-An Huang, Kay Chen Tan

## Abstract

Single-cell analysis has revolutionized our understanding of cellular heterogeneity, yet current approaches face challenges in efficiency and interpretability. In this study, we present scKAN, a framework that leverages Kolmogorov-Arnold Networks for interpretable single-cell analysis through three key innovations: efficient knowledge transfer from large language models through a lightweight distillation strategy; systematic identification of cell-type-specific functional gene sets through KAN’s learned activation curves; and precise marker gene discovery enabled by KAN’s importance scores with potential for drug repurposing applications. The model achieves superior performance on cell-type annotation with a 6.63% improvement in macro F1 score compared to state-of-the-art methods. Furthermore, scKAN’s learned activation curves and importance scores provide interpretable insights into cell-type-specific gene patterns, facilitating both gene set identification and marker gene discovery. We demonstrate the practical utility of scKAN through a case study on pancreatic ductal adenocarcinoma, where it successfully identified novel therapeutic targets and potential drug candidates, including Doconexent as a promising repurposing candidate. Molecular dynamics simulations further validated the stability of the predicted drug-target complexes. Our approach offers a comprehensive framework for bridging single-cell analysis with drug discovery, accelerating the translation of single-cell insights into therapeutic applications.

## Introduction

Single-cell technologies have revolutionized our understanding of biological processes and human diseases by offering unprecedented insights into cellular heterogeneity at high resolution^1,2^. A cornerstone of this revolution is cell-type annotation, which enables the identification of distinct cell populations across tissues, developmental stages, and organisms. This process provides a critical fundamental for understanding cellular functions and gene regulation in health and disease states^3–5^. The systematic identification of essential genes has provided critical insights into the molecular basis of biological processes^6^. However, how cell-essential genes differ across cell types remains poorly understood. These differentially essential genes likely encode tissue-specific modulators of key cellular functions and hold significant promise as therapeutic targets, particularly in cancer^7^.

Cell-type annotation methods rely on single-cell gene expression data to distinguish cell populations. Traditional methods like Seurat^8^, while widely used, often require extensive preprocessing steps and manual marker selection, limiting their scalability and automation^9^. To address these limitations, machine learning approaches have emerged and demonstrated significant progress. For example, CellTypist^10^ leveraged supervised learning techniques to identify 101 distinct cell types from over a million cells. Deep learning methods have further propelled the field. ACTINN^11^ employed neural networks to achieve rapid and accurate cell type identification. MetaTiME^12^ integrated single-cell gene expression to characterize meta-components, enabling the identification of heterogeneous cell types and states in tumor microenvironment. Meanwhile, Cellcano^13^ introduced a supervised learning framework that effectively mitigates the distributional shift between reference and target data, improving annotation accuracy.

Recently, Transformer-based large language models (LLMs) with attention mechanisms have shown immense promise for single-cell analysis. TOSICA^14^ pioneered one-shot annotation, adapting to new cell types with minimal training examples while maintaining interpretability through biologically understandable entities. scBERT^15^ introduced a BERT-inspired pretraining approach to capture gene-gene interactions from large-scale unlabeled single-cell RNA sequencing (scRNA-seq) data. Furthermore, LangCell^16^ was innovated by constructing unified representations of single-cell data and natural language during pretraining, enabling zero-shot cell identity understanding. Geneformer^17^ advanced context-aware attention-based learning with self-supervised pretraining on about 30 million single-cell transcriptomes. More recently, scGPT^18^, trained on over 33 million cells, established itself as a foundation model capable of diverse downstream applications, including cell-type annotation, multi-batch integration, and gene network inference. Meanwhile, GeneCompass^19^ further contributed by integrating prior biological knowledge across species. However, despite their potential, these LLM-based models often demand substantial computational resources for training, require frequent fine-tuning for new datasets, and struggle to provide cell-type-specific interpretability of gene functions and interactions^20^.

These limitations present three key technical challenges in single-cell analysis. First, despite extensive pretraining, these models require considerable fine-tuning to achieve acceptable accuracy on new data. Second, the adopted attention mechanisms focus on global context inherently limits their ability to identify cell-type specific genes. Third, current methods operate in isolation from downstream applications such as drug discovery, creating a gap between single-cell analysis and practical therapeutic development. These challenges call for a new paradigm that can naturally support the identification of cell-type specific genes and bridging the gap to drug repurposing.

To address these challenges in single-cell analysis, we propose a knowledge distillation-based framework called scKAN for single-cell analysis that leverages Kolmogorov-Arnold networks (KAN) architecture^21^. Unlike traditional neural networks with fixed activation functions and linear weight matrices, KAN implements learnable univariate functions as part of its architecture, rooted in the Kolmogorov-Arnold representation theorem. We chose KAN architecture for its demonstrated advantages in AI for Science applications^22^, particularly its inherent interpretability. Our key insight is that KAN’s trained activation curves and importance scores can provide cell-type-specific interpretability for gene selection, offering a natural framework for understanding cell-gene relationships. This architectural innovation enables scKAN to achieve three significant technical advancements. First, scKAN effectively transfers the pre-trained knowledge from LLMs into a lightweight framework through knowledge distillation, achieving efficient and accurate cell-type annotation without extensive fine-tuning. Second, the learned curves in scKAN naturally capture cell-gene representation patterns, enabling the identification of functionally similar cell-type-specific gene sets based on their expression characteristics. Third, the importance scores inherent to KAN’s architecture provide interpretable measures of gene importance, allowing for identifying cell-type-specific marker genes. This unified framework addresses the computational efficiency challenge and enables interpretable discovery of both gene sets and marker genes in a cell-type-specific manner.

Extensive experiments demonstrate that scKAN achieves competitive performance while maintaining computational efficiency. Our model effectively identifies cell types and their associated marker genes, revealing biologically meaningful relationships in complex cellular systems. Beyond technical validation, we successfully applied scKAN to pancreatic ductal adenocarcinoma (PDAC) research, where it identified critical gene signatures in pancreatic ductal cells.

Unlike traditional drug discovery approaches that rely solely on differential expression analysis, scKAN’s integration of cell-type-specific gene importance scores with activation curve patterns provides a novel framework for identifying druggable targets. Notably, scKAN enables the discovery of potential therapeutic targets that might be overlooked by conventional methods, particularly those with moderate expression levels but high functional significance. These findings led to the discovery of potential therapeutic targets, demonstrating scKAN’s value in translational research. Together, these results highlight scKAN’s capability to advance the methodological framework of single-cell analysis and its practical applications in disease research.

## Results

### Overview of the scKAN Framework

The workflow and model architecture of scKAN are shown in Fig. 1. scKAN aims to achieve accurate cell annotation and identify marker genes and gene sets for specific cell types. Fig. 1a provides a schematic overview of the biological context, highlighting the existence of diverse cell types within organisms and the role of single-cell technologies in generating gene expression matrices for individual cells. scKAN leverages these matrices to decipher cell types and identify marker genes and potential pathways.

**Figure 1.**
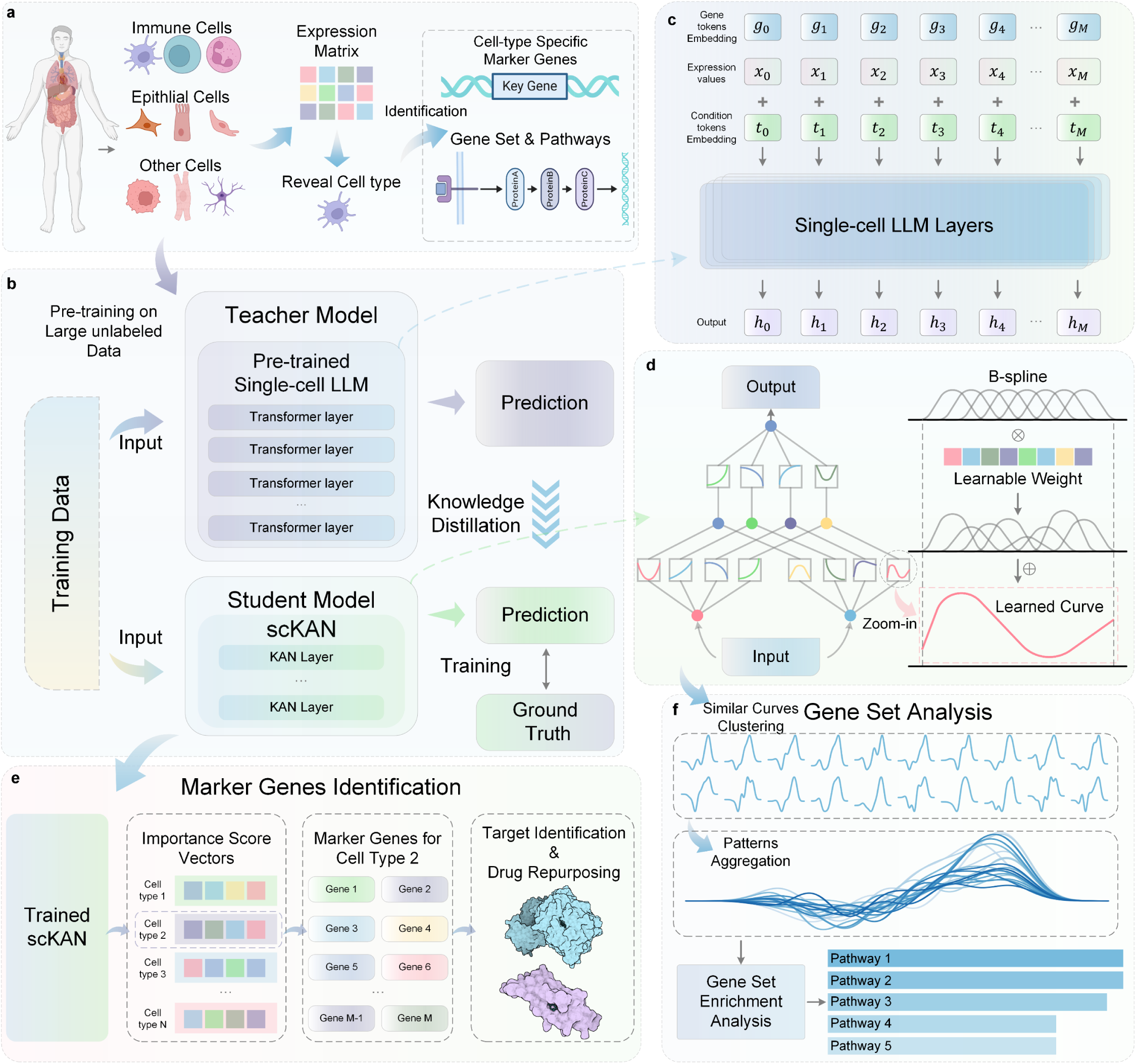
Overview of the scKAN framework for single-cell analysis and marker gene identification. **a,** Schematic illustration of the single-cell analysis workflow, showing the extraction of different cell types from biological samples, followed by generation of expression matrices and identification of cell-type-specific marker genes and pathways. **b,** Knowledge distillation framework architecture consisting of two main components: a teacher model (pre-trained single-cell LLM with multiple Transformer layers trained on large unlabeled datasets) and a student model (scKAN with multiple KAN layers). The framework enables knowledge transfer through prediction-based distillation while incorporating ground truth information during training. **c,** Structure of single-cell LLM layers, showing the integration of gene tokens, expression values, and condition embeddings to generate cell embeddings (output). **d,** Detailed neural network architecture of scKAN, illustrating the hierarchical structure of KAN layers where connections between nodes are represented by learnable B-spline activation functions instead of traditional weights. The right panel shows how these B-spline functions are learned and adjusted during training to capture complex relationships in the data. **e,** Marker gene identification process using trained scKAN, progressing from importance score vectors for different cell types to specific marker gene identification, ultimately supporting target identification and drug repurposing applications. **f,** Gene set analysis workflow showing the clustering of similar activation curves from trained scKAN, pattern aggregation, and subsequent pathway enrichment analysis to identify cell-type-specific pathways.

As illustrated in Fig. 1b-d, the core architecture of scKAN employs a knowledge distillation strategy. A pre-trained LLM based on the Transformer architecture^23^ serves as the teacher model, guiding a KAN-based module as the student model. The development of scKAN involves two steps: first, fine-tuning a LLM that has been pre-trained on large-scale unlabeled cellular data on specific datasets; second, training the KAN-based student model through knowledge distillation to enable it to integrate the teacher model’s prior knowledge with ground truth cell type information. In addition to the distillation process, we incorporate unsupervised learning objectives to enhance the discriminative power of learned feature representations^24^. Specifically, we employ a self-entropy loss^25^ to prevent over-concentration on dominant cell types and maintain sensitivity to rare cell populations. Furthermore, by leveraging Cauchy-Schwarz divergence^28^, a modified Deep Divergence-based Clustering (DDC) loss^26,27^ optimizes the relationships between hidden features and cluster assignements, ensuring alignment with ideal cell type distributions. This combined loss fuction, integrating distillation and unsupervised components improves the model’s generalization ability across different cell types.

To obtain a pre-trained LLM with extensive single-cell prior knowledge, we employ the state-of-the-art (SOTA) single-cell analysis foundation model, scGPT^18^, as the teacher model. This Transformer-based model has been extensively pre-trained on over 33 million cells, capturing representation patterns of various human cell types, including pancreatic and blood cells. As shown in Fig. 1c, it uses a gene encoder to encode gene id, applies binning to expression values to obtain expression embeddings, and incorporates condition embeddings for specific genes. By integrating these embedding inputs through multiple Transformer layers, this pretraining imbues the teacher model with a rich understanding of diverse human cell types for annotation.

We implement scKAN by comprising multiple KAN layers as the student model to learn from the teacher model while acquiring annotation capabilities on labeled cellular data. Following the Kolmogorov-Arnold representation theorem, the KAN model learns activation function curves for edges, rather than weights as in traditional multilayer perceptrons (MLPs). As shown in Fig. 1d, these curves, fitted using B-splines, capture underlying representation patterns, establishing latent representation connections between cells and genes.

After training, the edge scores in KAN, initially used for pruning, indicate the significance of activation function curves between nodes. In this study, these scores are hypothesized to reflect the importance of cell-gene relationships. As shown in Fig. 1e, these importance scores can identify marker genes for specific cell types, facilitating downstream analysis such as target identification of human diseases and functional characterization for specified cell types. Combined with drug-target affinity prediction methods, scKAN can potentially support drug repurposing studies.

Additionally, the similarity among learned activation function curves provides insights into gene co-expression patterns within specific cell types. As shown in Fig. 1f, clustering similar curves reveals functionally related gene sets. Subsequent gene set enrichment analysis can then pinpoint cell-type-specific pathways and uncover novel pathway-associated genes that have not been previously detected.

### scKAN Achieves Superior Performance in Cell-type Annotation

To evaluate the performance of scKAN in cell-type annotation tasks, we compared it with multiple baseline models: Geneformer^17^, Tosica^14^, and scGPT^18^, which are LLM-based single-cell foundation models; Cellcano^13^, a deep learning method employing knowledge distillation; Celltypist^10^, a machine learning-based method; and Seurat^8^, a classical single-cell analysis tool. Detailed information about these baseline methods is provided in the ‘Baseline’ subsection in the Methods section.

To ensure a fair comparison, we tested these methods on multiple datasets: PBMC^29^ dataset with 9,631 single cells across 19 cell types, Muto2021^30^ dataset containing more than 20,000 single cells, Human Pancreas Dataset^14^ (hPancreas) with 14,818 single cells, and Myeloid dataset^31^ (Mye) with 13,178 cells. These datasets include various cell types, such as immune and kidney cells. Detailed information about these datasets is provided in the ‘Dataset’ subsection in the Methods section.

Fig. 2 shows the performance comparison between scKAN and baseline methods across these datasets. We used classical multiple classification metrics including accuracy, macro precision, macro recall, and macro F1 score, to assess model performance comprehensively.

**Figure 2.**
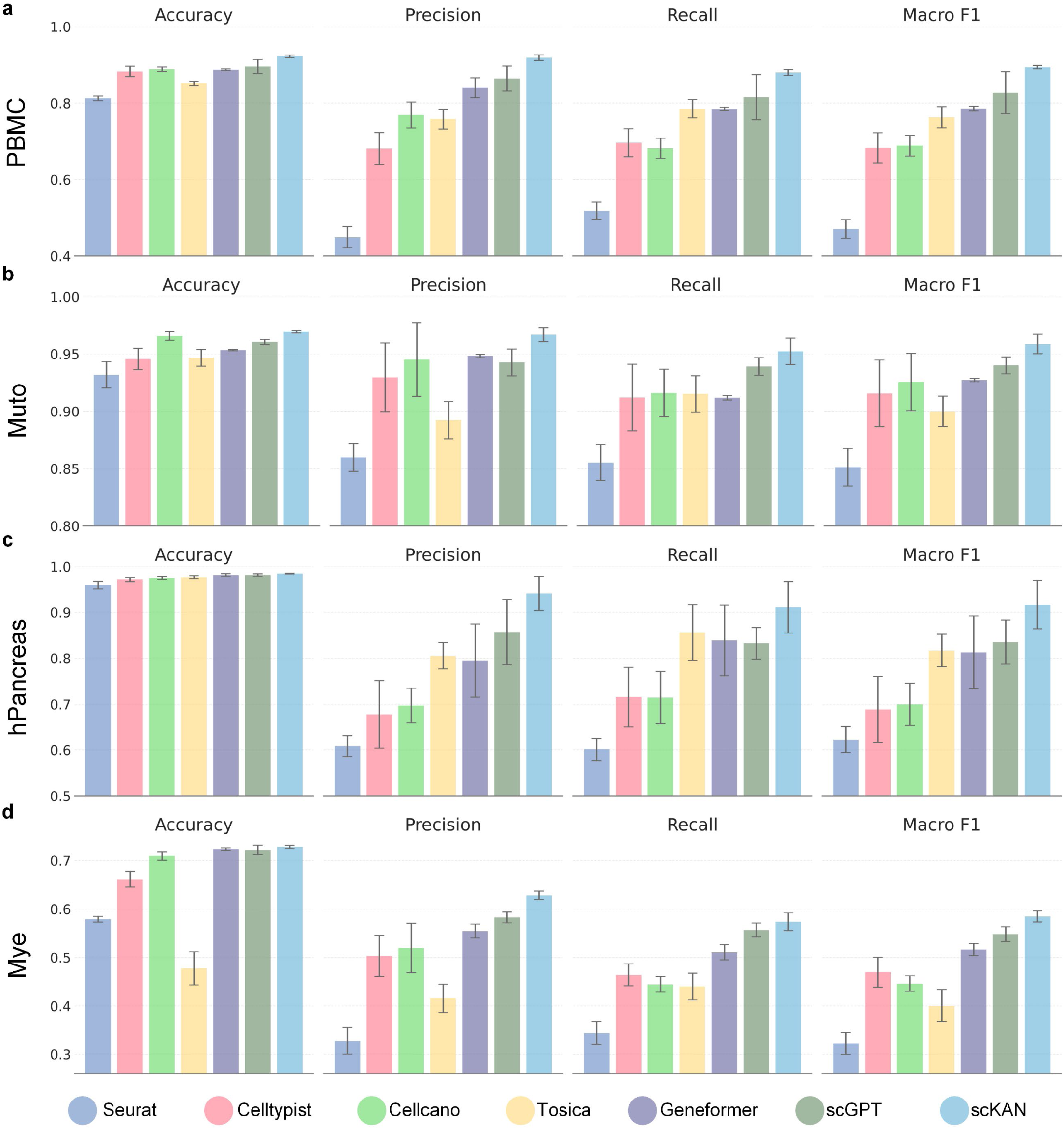
Comparative performance evaluation of scKAN against baseline methods across multiple datasets using 5-fold cross-validation. **a-d,** Comparison of test set performance in cell-type annotation across four datasets (PBMC, Muto, hPancreas, and Mye) using four metrics: accuracy, macro precision, macro recall, and macro F1 score. Results are averaged across five test sets from 5-fold cross-validation, with error bars representing standard deviation. Baseline methods include traditional tools (Seurat), machine learning approaches (Celltypist), deep learning methods (Cellcano), and foundation single-cell analysis large language models (Tosica, Geneformer, and scGPT). scKAN consistently outperforms baseline methods across different datasets and metrics, with significant improvements in accuracy and macro F1 scores.

scKAN consistently outperformed all baseline methods across all datasets and metrics, including the single-cell foundation models such as scGPT, Geneformer, and Tosica. This demonstrates the superior performance of scKAN while also underscoring the limitations of current single-cell foundation models, which still face challenges in cell-type annotation despite their versatility. Specifically, the experiment results are shown in Fig. 2, Supplementary Fig. S1, and Table S1. On average, scKAN achieved a 1.06% improvement in accuracy and a 6.63% improvement in macro F1 score compared to the second-best models across all datasets. Notably, on the PBMC dataset, scKAN achieved its highest accuracy improvement of 2.95% over the second-best model, scGPT. For the hPancreas dataset, scKAN demonstrated its most significant macro F1 score improvement, with a 9.77% increase over scGPT, reaching a score of 0.917. These consistent improvements across different datasets demonstrate that scKAN is a reliable and effective tool for cell-type annotation tasks.

Statistical analysis further validated these results. Using t-tests, we found significant improvements in the comprehensive metrics of accuracy and macro F1 score. For the PBMC dataset, both macro F1 and accuracy showed significant improvements (p < 0.05) compared with the second-best methods. In the Muto2021 dataset, while accuracy differences were insignificant, the macro F1 score showed significant improvement (p < 0.05). The hPancreas dataset showed significant improvements in both accuracy and macro F1 (p < 0.05), and the Mye dataset showed significant improvement in macro F1 score (p < 0.01) with no significant difference in accuracy. These statistical results further confirm that scKAN provides significantly better and more stable performance in cell-type annotation than existing methods.

Beyond performance advantages, scKAN demonstrates remarkable computational efficiency. When evaluated on the Muto2021 dataset, scKAN achieves a 15.8-fold reduction in GPU memory consumption and 5.4-fold faster training speed compared to scGPT, while requiring only 2.3% of the parameters (Table S2). This substantial improvement in computational efficiency makes scKAN more accessible for widespread adoption in the research community.

Additionally, we provided a detailed visualization analysis to further demonstrate scKAN’s performance. As illustrated in Fig. S1, the confusion matrices and UMAP visualizations across four datasets demonstrate scKAN’s consistent and accurate performance across different cell types. In the PBMC dataset, the confusion matrix shows high diagonal values, indicating accurate classification across all 19 cell types, with robust performance in identifying major immune cell populations. The UMAP visualization of PBMC data shows that the predicted cell type distributions closely match the annotated labels, with clear boundaries between different cell populations. Similar high-quality results were observed in the Muto2021 dataset, where the model accurately distinguished between different cell types, including CNT, DCT, ENDO, and LEUK populations. For the hPancreas dataset, scKAN successfully identified pancreatic cell types, including alpha, beta, and ductal cells, with the UMAP plot showing nearly identical clustering patterns between predicted and annotated labels. In the more challenging Myeloid dataset with its complex hierarchy of immune cell subtypes, scKAN maintained reliable performance, accurately distinguishing between closely related myeloid cell populations as shown in both the confusion matrix and UMAP visualization. These visual results comprehensively demonstrate scKAN’s robust capability in cell-type annotation across diverse tissue types and cellular compositions. Overall, these detailed comparisons and analyses establish scKAN as a reliable and superior tool for single-cell annotation tasks.

### scKAN Maintains Robust Performance in Cross-study and Cross-disease Settings

To evaluate scKAN’s performance under more realistic conditions, we conducted cross-study experiments on the hPancreas dataset. Following Tosica’s experimental setup^14^, we divided the hPancreas dataset into non-overlapping reference and query sets from completely independent studies. The reference set contained 10,600 single-cell samples from GSE84133 and GSE85241, covering 14 cell types, while the query set included 4,218 previously unseen single-cell samples from three distinct studies (GSE81608, E-MTAB-5061, and GSE86473), representing 11 cell types. This setup ensures a strict evaluation of the model’s generalization ability across independent datasets with no sample overlap.

As shown in Fig. 3 and Supplementary Table S3, in this cross-study setting, scKAN achieved 97.42% accuracy and a macro F1 score of 0.7342, showing improvements of 1.01% and 2.03% over the second-best models Tosica and scGPT, respectively. As shown in Fig. 3a-d, the pie charts clearly illustrate scKAN’s superior performance across all evaluation metrics compared to other methods. Even under this more challenging cross-study scenario, scKAN maintained high performance with accuracy close to 98%, demonstrating strong generalization ability. The confusion matrix (Fig. 3e) reveals high diagonal values for most cell types, indicating accurate classification performance. The inability to effectively annotate MHC class II cells was also observed across all baseline models due to this severe data imbalance. The UMAP visualization (Fig. 3f) further supports these findings, showing consistent cell type distributions between reference and query sets, with predicted labels closely matching the actual cell types.

**Figure 3.**
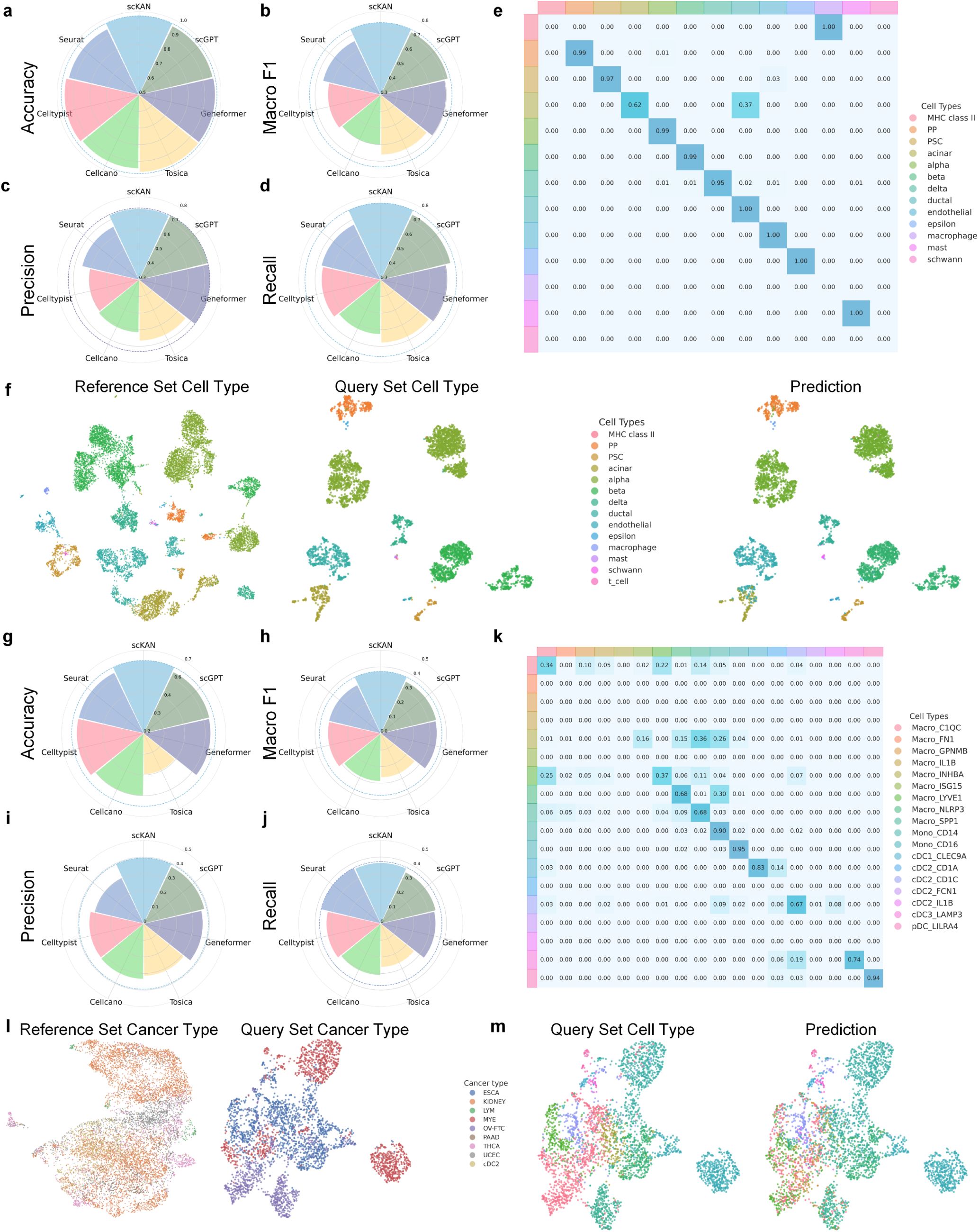
Performance evaluation of scKAN in cross-study and cross-disease settings. **a-d,** Comparative performance metrics (accuracy, macro F1, precision, and recall) across different methods in the cross-study pancreas dataset experiment. The pie charts show scKAN’s superior performance compared to baseline methods. **e,** Confusion matrix showing cell type classification performance across 13 pancreatic cell types in the cross-study setting. **f,** UMAP visualization of pancreas data showing reference set cell types (left), query set cell types (middle), and scKAN predictions (right), demonstrating consistent cell type distribution patterns. **g-j,** Performance metrics displayed as pie charts for the cross-disease myeloid dataset experiment, showing scKAN’s maintained advantage in this challenging setting. **k,** Confusion matrix demonstrating classification performance across 17 myeloid cell types in the cross-disease setting. **l,** UMAP visualization comparing reference set (left) and query set (right) cancer type distributions. **m,** UMAP visualization of query set cell types (left) and corresponding scKAN predictions (right), showing preserved cell type distribution patterns despite the cross-disease setting.

We further tested scKAN under a more challenging cross-disease setting using the Mye dataset. As illustrated in Fig. 3l-m, we selected six cancer types for the reference set (9,748 cells) and three for the query set (3,430 cells). The pie charts (Fig. 3g-j) demonstrate that in this cross-disease scenario, scKAN achieved an accuracy of 63.84% and a macro F1 score of 0.3726, surpassing the second-best model scGPT by 4.48% and 7.44%, respectively. Although overall performance decreased in this more challenging setting, scKAN maintained its advantage over baseline methods across all metrics, including macro F1 score.

The confusion matrix (Fig. 3k) shows the classification results across different cell types, revealing a block-diagonal pattern that indicates strong classification performance within related cell type groups. The model achieved high accuracy (>0.6) for most cell types, with lower performance only in C1QC, INHBA, and LYVE1 classifications, similar to the hPancreas dataset, primarily due to limited sample sizes for these cell types. The UMAP visualizations (Fig. 3l-m) demonstrate that despite the challenging cross-disease setting, scKAN successfully maintained the overall structure of cell type distributions between reference and query sets, with predicted labels showing strong concordance with actual cell types. These cross-study and cross-disease results demonstrate that scKAN can effectively learn intrinsic patterns in single-cell data, enabling accurate cell-type annotation across different experimental conditions.

### Ablation Studies Validate the Essential Components of scKAN

To systematically evaluate the contribution of each component in scKAN, we conducted comprehensive ablation studies by creating three model variants: ‘W/o Teacher’ removes the teacher model to assess the impact of knowledge distillation, ‘W/o Cluster’ eliminates the clustering loss to evaluate the importance of maintaining cell-type-specific feature representations, and ‘Replaced by MLP’ substitutes the KAN module with a MLP architecture to examine the advantages of our KAN-based design. These variants were designed to validate our architectural choices and understand the role of each component in achieving optimal performance.

We evaluated these variants using accuracy and macro F1 scores across all four datasets. These two metrics were chosen as they provide complementary insights: accuracy reflects the overall classification performance, while macro F1 score better captures the model’s performance on imbalanced cell populations. We omitted individual precision and recall metrics to focus on their combined effects through the F1 score, providing a more concise yet comprehensive evaluation.

The ablation results in Table 1 demonstrate the importance of each component. ‘W/o Teacher’ led to consistent performance drops across all datasets, particularly noticeable decreases in macro F1 scores. On the PBMC dataset, the macro F1 score dropped from 0.894 to 0.869, while on hPancreas, it decreased from 0.917 to 0.871, confirming the value of knowledge transfer from the pre-trained language model. Similarly, ‘W/o Cluster’ reduced performance, especially in the macro F1 scores of more challenging datasets. For instance, on the hPancreas dataset, removing the clustering loss led to a decrease in the macro F1 score from 0.917 to 0.892, highlighting its role in maintaining robust cell-type representations. When replacing KAN with MLP, we observed an interesting trade-off. The MLP variant achieved higher accuracy on the Mye dataset, reaching 0.747 compared to scKAN’s 0.728. However, it showed consistently lower macro F1 scores across datasets. This discrepancy likely stems from MLP’s tendency to optimize for majority cell types at the expense of rare populations, particularly evident in the complex and imbalanced Mye dataset. Moreover, replacing KAN with MLP eliminates the interpretability advantages provided by KAN’s learned activation curves and importance scores, losing the ability to gain biological insights from the model’s decision-making process.

**Table 1.**
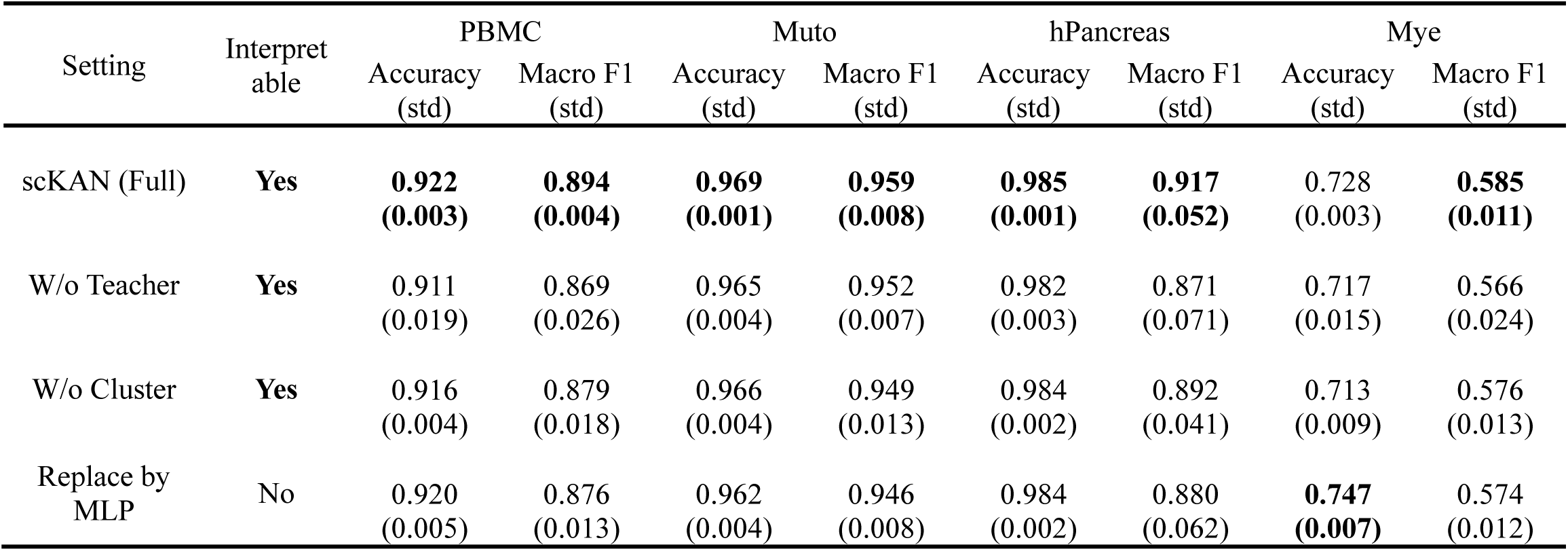
Ablation study evaluating the contribution of key components in scKAN. The performance metrics include accuracy and macro F1 scores with standard deviations (shown in parentheses) across four datasets: PBMC, Muto, hPancreas, and Mye. ‘W/o Teacher’ represents the model without knowledge distillation, ‘W/o Cluster’ removes the clustering-based loss functions, and ‘Replace by MLP’ substitutes the KAN module with a traditional multilayer perceptron. Bold numbers indicate the best performance. The ‘Interpretable’ column indicates whether the model variant maintains the inherent ability to provide interpretable insights into cell-gene relationships.

These ablation studies demonstrate that each component plays a crucial role in scKAN’s overall performance and functionality. The knowledge distillation framework provides essential prior knowledge, the clustering loss helps maintain robust cell-type-specific features, and the KAN architecture offers a unique combination of competitive performance and biological interpretability. Alternative architectures like MLP might achieve comparable accuracy in specific scenarios. However, our design choices offer comprehensive benefits. These include balanced performance across cell types and enhanced interpretability. These advantages strongly support the current architecture of scKAN.

### scKAN Enables Accurate Gene Set Identification and Pathway Analysis

After validating the model’s cell annotation capabilities, we investigated its ability to identify biologically meaningful gene sets. scKAN learns gene characteristics across different cell types by distilling knowledge from single-cell LLMs, implicitly encoding cell type-level gene expression patterns within gene-cell activation curves, thereby enabling gene set identification.

First, as shown in Fig. 4a, we visualized gene programs clustered by these activation curves from scKAN in the PBMC dataset, revealing diverse expression patterns across cell types. The model-identified gene programs indicated that the model captures more than just simple, single-pattern programs. Furthermore, Fig. 4b demonstrates that genes with similar scKAN curves within gene programs showed remarkably high similarity, particularly when contrasted with the disorder of randomly selected activation curves shown in Fig. 4c.

**Figure 4.**
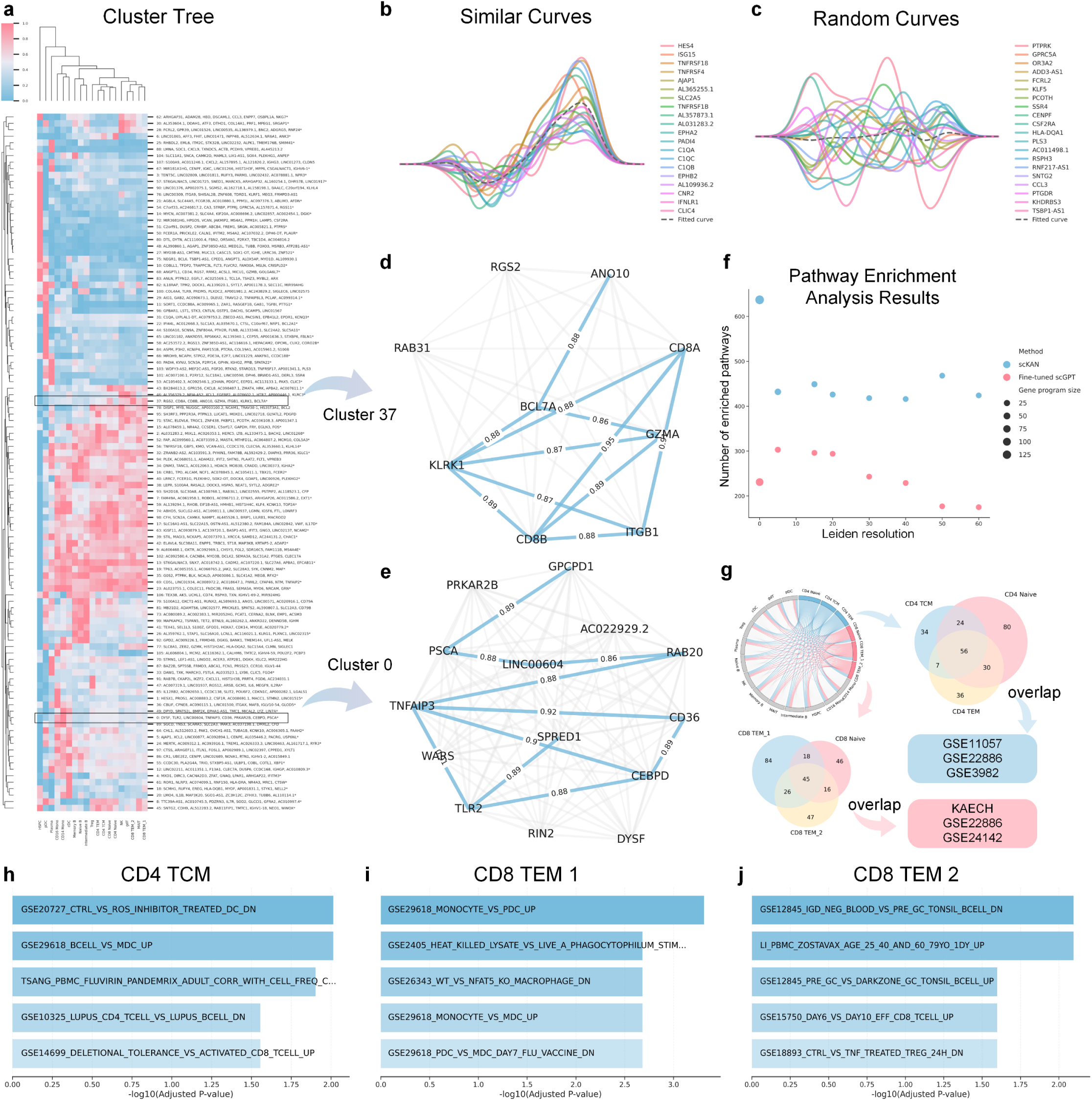
Gene set identification and pathway analysis capabilities of scKAN. **a,** Hierarchical clustering of gene expression patterns derived from scKAN activation curves, with heatmap showing expression patterns across different cell types. **b,** Example of similar activation curves from a functionally related gene cluster, demonstrating coherent patterns. **c,** Random gene activation curves are shown for comparison, illustrating the contrast with functionally related curves. Gray dashed lines in (b) and (c) show the mean fitted curves. **d-e,** Network visualization of two representative gene clusters: (d) T cell-related gene cluster (Cluster 37) and (e) inflammation-related gene cluster (Cluster 0), with edge weights indicating similarity scores between gene activation curves. **f,** Comparison of pathway enrichment analysis results between scKAN and scGPT across different Leiden resolutions, showing the number of enriched pathways (dot size indicates gene program size). **g,** Venn diagram and circular visualization showing pathway overlap between CD4 and CD8 T cell populations, with data source labels (GSE11057, GSE3982, GSE22886, KAECH) indicated. **h-j,** Top 5 enriched pathways (-log10 adjusted p-value) for specific T cell populations: (h) CD4 TCM, (i) CD8 TEM 1, and (j) CD8 TEM 2, demonstrating cell-type-specific pathway enrichment.

Specifically, the gene programs encompassed various functionally distinct programs. For instance, as illustrated in Fig. 4d, Cluster 37 contained several T cell-related genes. These included the CD8A and CD8B genes, which work together to form the CD8 complex essential for T cell recognition^32^. The cluster also contained GZMA, a crucial effector molecule in CD8+ T cells^33^, along with KLRK1, which is an activation receptor in both NK and T cells^34^. Additionally, ITGB1 was identified as playing a vital role in T cell adhesion and migration processes^35,36^. The signaling-related components of this cluster included RGS2, which regulates G-protein signaling, and RAB31, which mediates intracellular transport^37,38^. Notably, the activation curves of these functionally related genes exhibited high similarity.

Similarly, Cluster 0, shown in Fig. 4e, revealed a distinct group of genes involved in inflammation and immune response pathways. TLR2 emerged as a key component, serving as a critical receptor for pathogen recognition and initiation of innate immunity^39^. The cluster also included TNFAIP3, a central regulator of inflammatory responses, alongside CD36, which functions as an important immune cell surface receptor^40^. CEBPD was also identified within this cluster, playing a crucial role in the transcriptional regulation of inflammatory genes^41^. The functional coherence of this cluster was further supported by the high similarity scores (>0.88) among the scKAN-derived curves for these genes, providing additional validation of the model’s effectiveness.

To further assess scKAN’s comprehension of gene expression patterns, we compared it with scGPT^18^, a SOTA single-cell LLM. We compared the number of unique pathways identified through Gene Set Enrichment Analysis (GSEA) using immune pathways from molecular signatures databases^42^ across different Leiden resolution levels. As shown in Fig. 4f, scKAN consistently captured more unique pathways across various resolutions. Notably, scKAN’s ability to cluster across different cell types, unlike scGPT embeddings, resulted in a greater number of clusters. While this numerical comparison alone is not definitive, it highlights scKAN’s advantage in cell-type-specific pathway identification.

Further pathway analysis, as illustrated in Fig. 4g, revealed overlapping pathways identified by scKAN across different cell types. Within the CD4 cell populations (CD4 TCM, CD4 Naïve, and CD4 TEM), we identified several significantly enriched pathways characteristic of T cell development and function, with 56 pathways shared among all three subsets. Notably, pathways from the GSE11057 collection distinguished memory CD4+ T cells from peripheral blood mononuclear cells, capturing memory T cell states with characteristic gene expression patterns^43^. We also identified GSE22886 pathways^44^ revealing molecular distinctions between naïve CD4+ T cells and monocytes, while GSE3982 pathways^45^ highlighted the unique molecular features of central memory CD4+ T cells through comparison with B cells.

Similarly, within CD8 cell populations (CD8 TEM_1, TEM_2, and naïve), we observed enrichment in pathways associated with CD8+ T cell differentiation and activation, with 45 pathways shared among all three populations. The KAECH pathways^46^ were particularly informative, capturing the dynamic transcriptional changes during CD8+ T cell activation and the transition from naïve to effector memory states. GSE22886 pathways^44^ highlighted CD8+ T cell-specific molecular signatures through comparisons with B cells and monocytes, while GSE24142 pathways^47^ revealed programs involved in early T cell development. All enriched pathways and their biological implications are provided in Supplementary Table S4.

Finally, as demonstrated in Fig. 4h-j, Supplementary Fig. S2, and Table S5, the top enriched pathways identified by scKAN curves for specific cell types are well-supported by previous literature. For CD4 TCM, the top enriched pathway reveals its critical role in modulating DC function, where CD4 TCM cells may influence DC antigen presentation through cytokine-mediated regulation of oxidative stress responses^48^. CD8 TEM1-associated pathway (GSE29618_MONOCYTE_VS_PDC_UP) demonstrates its involvement in regulating both monocyte and plasmacytoid dendritic cell functions, potentially impacting their antigen presentation capabilities through cytokine-mediated activation^49^. The CD8 TEM2-enriched first-rank pathway indicates its role in modulating B cell activation states and immunoglobulin gene expression, particularly in the context of germinal center B cell development^50^. Additional cell-type-specific pathways were also identified, as shown in Supplementary Fig. S2: NK cells exhibited enrichment in BCG vaccine-induced bactericidal activity pathways^51^, while naive B cells showed association with vaccine-induced PBMC responses^52^, and cDCs demonstrated enrichment in pathways related to vaccine-mediated immune responses^53^. Collectively, these comprehensive analyses validate scKAN’s effectiveness in interpretable gene set identification and its capability for cell-type-specific characterization.

### scKAN Discovers Reliable Cell-type-specific Marker Genes

Identifying cell-type-specific marker genes is crucial for understanding cellular identity and function. Here, we demonstrate that scKAN not only enables interpretable gene set identification but also provides a novel approach for marker gene discovery through interpretable importance scores. To systematically evaluate scKAN’s capabilities in marker gene identification, we conducted comprehensive analyses across multiple dimensions using both computational and biological validation approaches.

We performed differential expression analysis on the PBMC dataset encompassing 19 distinct cell types. We leveraged a well-trained scKAN model for each cell type to identify the top 20 candidate marker genes and evaluated their expression patterns using Wilcoxon rank-sum tests. Fig. 5a presents a comprehensive visualization where dot size represents the magnitude of expression fold change, color indicates regulatory direction, and black borders denotes statistical significance (adjusted p-value < 0.05). The analysis revealed that scKAN’s top 10 marker genes generally exhibited more pronounced differential expression patterns. However, some cell types, such as HSPCs, showed less significant differentiation. Markers ranked 10-20 displayed more varied patterns, including genes with modest expression differences. This dual capability of identifying highly expressed and subtly regulated marker genes provides opportunities to discover novel regulatory mechanisms in cell-type-specific gene expression.

**Figure 5.**
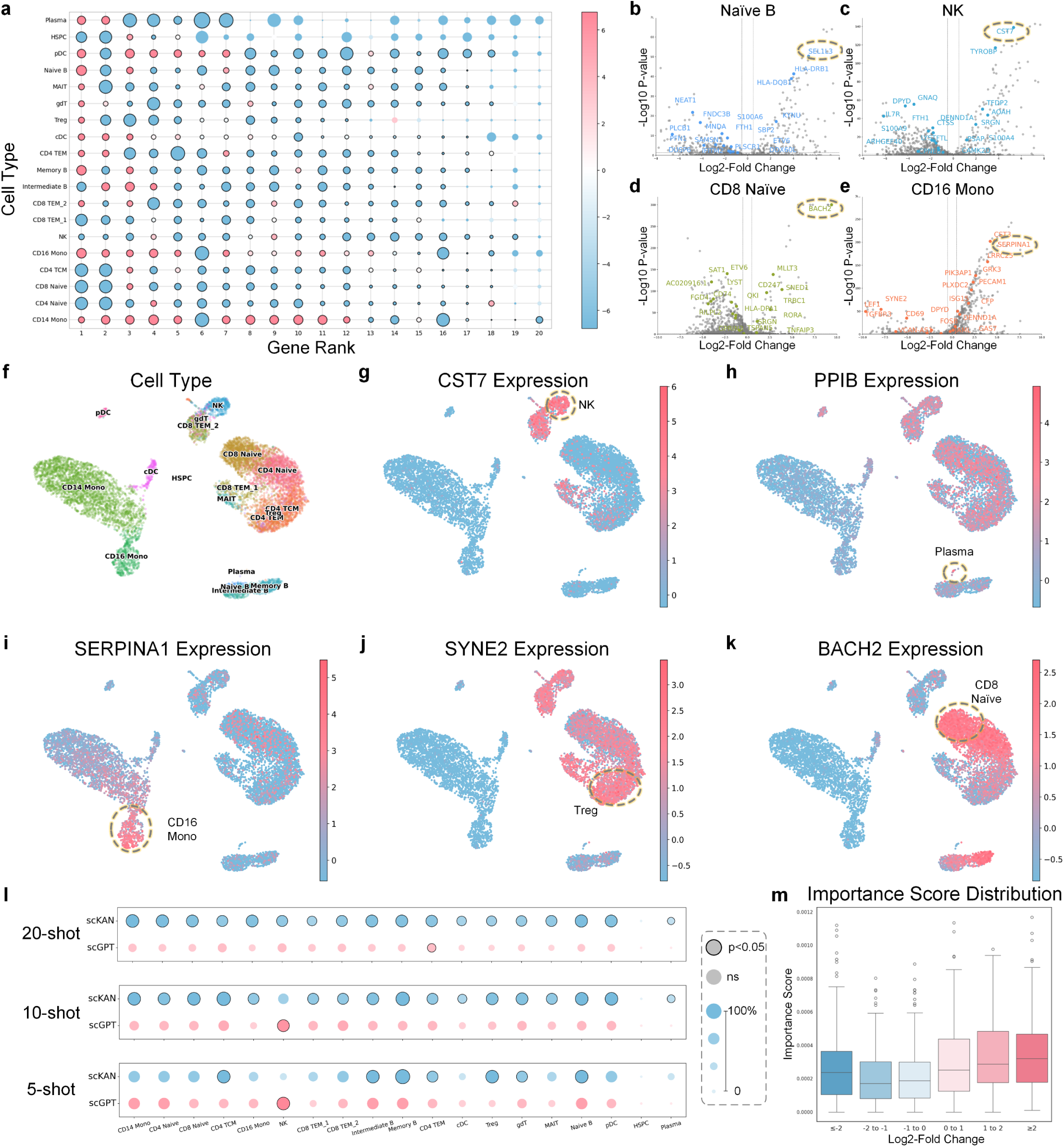
Marker gene identification and validation using scKAN. **a,** Dot plot showing expression patterns of top 20 marker genes identified by scKAN across 19 cell types. Dot size indicates expression fold change magnitude, color represents regulatory direction (-log10 p-value), and black borders denote statistical significance (adjusted p-value < 0.05). **b-e,** Volcano plots highlight differentially expressed genes in selected cell populations: (b) Naïve B cells, (c) NK cells, (d) CD8 Naïve cells, and (e) CD16 Mono cells, with marker genes highlighted in dashed circles. **f,** UMAP visualization of cell type distributions in the PBMC dataset. **g-k,** Expression patterns of validated marker genes shown on UMAP plots: CST7 (g), PPIB (h), SERPINA1 (i), SYNE2 (j), and BACH2 (k), with color intensity indicating expression level and dashed circles highlighting cell-type-specific expression. **l,** Comparative analysis of marker gene identification performance between scKAN and scGPT under different few-shot settings (20-shot, 10-shot, 5-shot), with dot size indicating the proportion of differentially expressed genes and black borders showing statistical significance (p<0.05). **m,** The distribution of scKAN importance scores across different log2-fold change ranges, showing the relationship between importance scores and expression differences.

As shown in Fig. 5b-d and Supplementary Fig. S3, detailed examination through volcano plots further validates scKAN’s capacity to identify both strongly and subtly differentially expressed genes. The biological relevance of scKAN-identified markers was corroborated by extensive literature support. Key examples include SEL1L3 in naïve B cells, which has been previously established as a crucial regulator of B cell development^54^; CST7 in NK cells, known for its role in cytotoxic functions^54^; BACH2 in CD8 naïve cells, essential for T cell homeostasis^55^; SERPINA1 in CD16 monocytes, involved in inflammatory responses^56,57^; and PPIB in plasma cells, critical for antibody production^54^. UMAP visualizations (Fig. 5f-m and Supplementary Fig. S4) demonstrate the distinct spatial distribution of these markers, confirming their cell-type-specific expression patterns.

To benchmark scKAN’s interpretable marker gene identification capabilities, we conducted a systematic comparison with scGPT, a SOTA single-cell foundation model, under 20-shot, 10- shot, and 5-shot settings. We evaluated the top 20, top 10, and top 5 potential marker genes for each cell type. Fig. 5n illustrates the proportion of differentially expressed genes captured by each model across different cell types, with black-bordered circles indicating statistical significance determined by hypergeometric testing. scKAN demonstrated superior performance in capturing differentially expressed genes across most cell types, significantly enriching the overall differentially expressed gene pool. The only exception was NK cells under 10-shot and 20-shot conditions, where scGPT showed comparable performance. These statistical results strongly support scKAN’s effectiveness in capturing cell-type-specific gene expression patterns.

Furthermore, Fig. 5o displays the distribution of scKAN-derived importance scores across different log2-fold change ranges. While the distributions showed subtle variations across fold change intervals, a trend emerged where higher absolute log2-fold changes corresponded to elevated importance scores. This pattern suggests that scKAN’s importance metric captures not only expression magnitude but also more complex regulatory relationships, providing a more comprehensive framework for marker gene identification than traditional differential expression analysis alone.

These comprehensive analyses demonstrate that scKAN provides a robust and interpretable approach for marker gene identification, capturing both strong and subtle cell-type-specific signals. The model’s effectiveness, validated through statistical and biological approaches, establishes its utility for marker gene discovery in single-cell analysis.

### scKAN Enables Systematic Drug Repurposing for PDAC Treatment

PDAC remains one of the most lethal malignancies, with a dismal five-year survival rate below 8.5%^58^. Despite extensive research efforts, therapeutic options remain limited, highlighting the urgent need for novel drug development approaches^59^. This urgent clinical need motivated us to explore scKAN’s potential in interpretable drug discovery through a systematic drug repurposing study targeting PDAC.

We designed a comprehensive workflow to drug repurposing for PDAC. Initially, we employed scKAN trained on the hPancreas dataset to identify ductal cell-specific marker genes. Fig. 6a highlights these markers in the volcano plot, showing diverse differential expression patterns consistent with our previous analyses. Notably, literature validation revealed that 12 out of the top 20 marker genes had been previously implicated in pancreatic cancer or specifically PDAC^60–63^ (Fig. 6b). Among the top 10 genes, 8 were validated, representing a significantly high validation rate ^64–71^, further underscoring scKAN’s reliability in identifying cell-type-specific markers. Detailed information about supporting work for those potential genes is shown in Supplementary Table S6.

**Figure 6.**
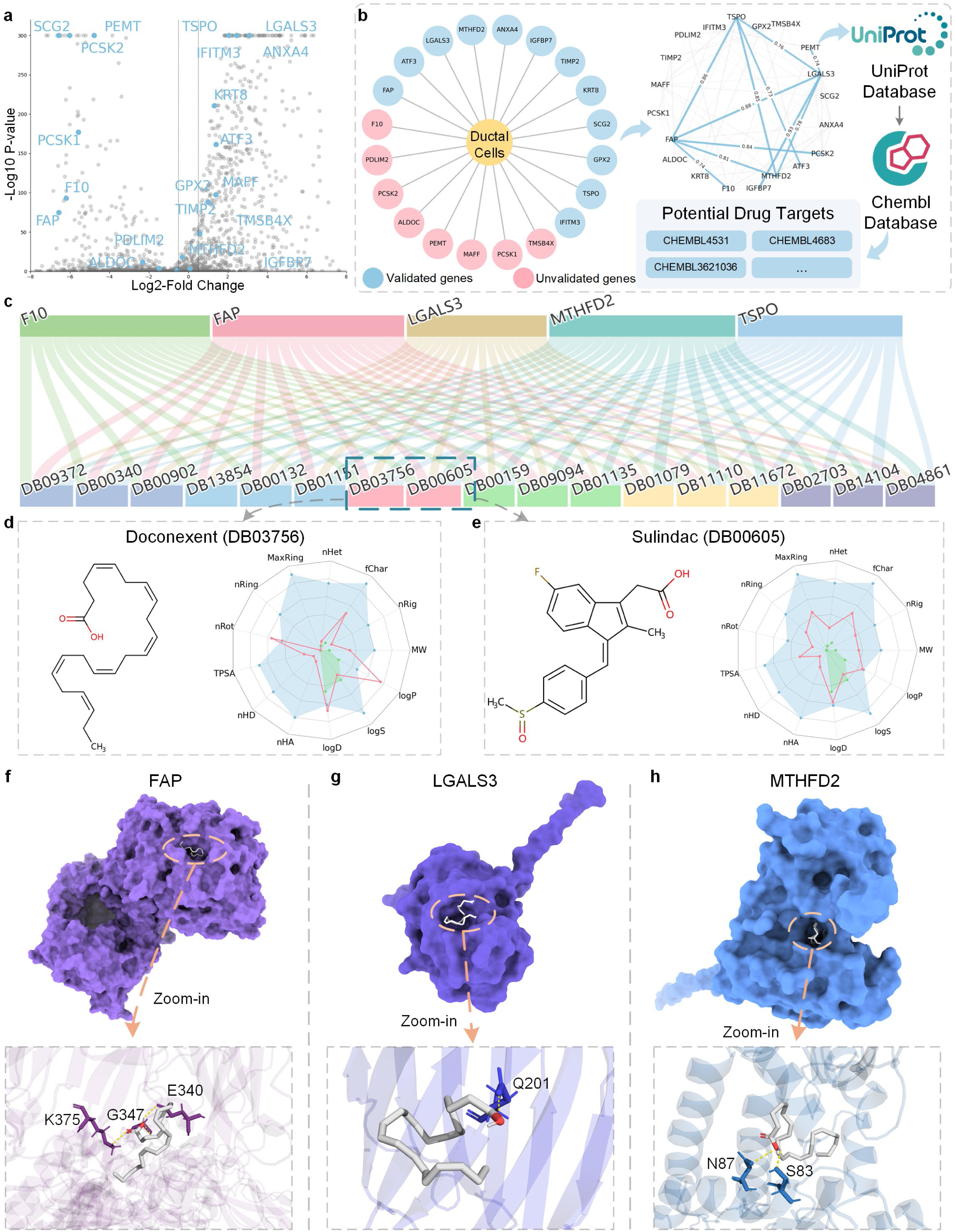
Drug repurposing analysis for PDAC treatment using scKAN and molecular verification. **a,** Volcano plot showing differentially expressed genes in ductal cells. Potential marker genes identified by scKAN are highlighted in blue. **b,** Network showing scKAN-identified marker genes for ductal cells, with previously validated genes in blue and unvalidated genes in pink. Similarity graph constructed from scKAN activation curves showing relationships between those marker genes, followed by potential drug target identification through UniProt and drug-target affinity data retrieval from the ChEMBL database. **c,** Sankey diagram showing predicted binding affinity relationships between potential protein targets and FDA-approved drugs, with top candidates DB03756 (Doconexent) and DB00605 (Sulindac) highlighted in pink. **d-e,** 2D structures and ADMET radar plots for top drug candidates: (d) Doconexent (DB03756) and (e) Sulindac (DB00605). **f-h,** Molecular docking results showing binding poses of Doconexent with three target proteins: (f) FAP, (g) LGALS3, and (h) MTHFD2, with surface representations (top) and detailed binding pocket views (bottom).

To further explore the functional relationships among these potential markers, we leveraged scKAN’s activation curves to construct a similarity graph for analyzing gene-gene associations among these marker genes. This analysis enabled us to identify highly interconnected gene modules, from which we extracted a tightly correlated gene set comprising FAP, F10, IGFBP7, MTHFD2, ATF3, PCSK2, LGALS3, PEMT, and TSPO. We then mapped these genes to their corresponding protein targets using the UniProt database^72^ and searched for existing drug-target interaction data in the ChEMBL database^73^. This systematic database integration revealed that only five targets - F10, FAP, LGALS3, MTHFD2, and TSPO - had corresponding drug-target datasets available. Subsequently, we employed the NHGNN^74^, a widely used SOTA drug-target affinity prediction and drug repurposing method, fine-tuning it on our collected datasets. Screening 2,509 FDA-approved drugs against these targets (Fig. 6c) identified Doconexent (DB03756) and Sulindac (DB00605) as the top two candidates, ranking first and second, respectively, based on combined prediction scores. Notably, Sulindac has been previously demonstrated to have therapeutic potential in pancreatic cancer treatment^75^, providing external validation for our computational approach and supporting scKAN’s utility in identifying clinically relevant drug targets.

To evaluate our identified candidate’s drug-likeness and potential pharmacological properties, we employed ADMETLab 3.0^76^ to assess their absorption, distribution, metabolism, excretion, and toxicity (ADMET) properties. The 2D structural visualization and ADMET analysis results are shown in Fig. 6d-e, revealing that both molecules mostly satisfy ADMET criteria, falling within acceptable ranges for most parameters. Only Doconexent showed elevated nRot, LogD, and LogP values, while both molecules exhibited lower than optimal LogS values, suggesting potential for further molecular optimization while maintaining drug-like properties.

To further investigate Doconexent, our top-ranked candidate that outperformed the literature-supported Sulindac in virtual screening, we sought to characterize its potential binding mechanisms through molecular docking. We employed CB-Dock2^77^, a cavity-detection-guided docking approach, to examine the binding interactions between Doconexent and PDAC’s potential protein targets. Crystal structures for these targets were obtained from the RCSB Protein Data Bank^78^ for FAP (PDB ID: 1Z68^79^), LGALS3 (PDB ID: 1KJL^80^), MTHFD2 (PDB ID: 5TC4^81^), and F10 (PDB ID: 4Y6D^82^). For TSPO, due to the absence of experimental structures, we generated a predicted structure using AlphaFold3^83^ (AF3) web server.

The molecular docking analysis yielded remarkably high CB-Dock scores (>6) across all protein targets, indicating strong binding potential. The docked complexes and detailed binding interfaces are visualized in Fig. 6f-h and Supplementary Fig. S5, where the upper panels show the surface representation of protein-ligand complexes, and the lower panels present detailed views of the binding pockets. Multiple hydrogen bond formations were observed within these binding pockets, further supporting the stability of these protein-ligand interactions. These structural analyses provide molecular-level evidence supporting Doconexent’s potential therapeutic application in PDAC.

This comprehensive drug repurposing workflow for PDAC demonstrates scKAN’s potential as a computational strategy for drug target discovery, providing a foundation for future experimental validation studies.

### scKAN-identified Drug Candidate Shows Stable Binding to potential Targets

Following our molecular docking analyses, we extended our investigation through molecular dynamics (MD) simulations to evaluate the binding persistence and stability of the predicted Doconexent-target complexes. These simulations were designed to assess the maintenance of key protein-ligand interactions under physiological conditions, providing insights into the stability of binding modes identified from molecular docking. We performed 100 ns all-atom molecular dynamics simulations at 310.15 K and 1 atm pressure, with detailed protocols in the Methods section.

The initial (gray) and final (blue) conformations of the five protein-ligand complexes are shown in Fig. 7 (left panels). Structural comparison before and after simulation revealed high overall conformational conservation, with major variations primarily observed in the terminal residues of the target proteins (highlighted by orange dashed circles). Notably, Doconexent maintained its position within the initial binding pockets across all five complexes throughout the 100 ns simulations, suggesting stable and effective protein-ligand interactions.

**Figure 7.**
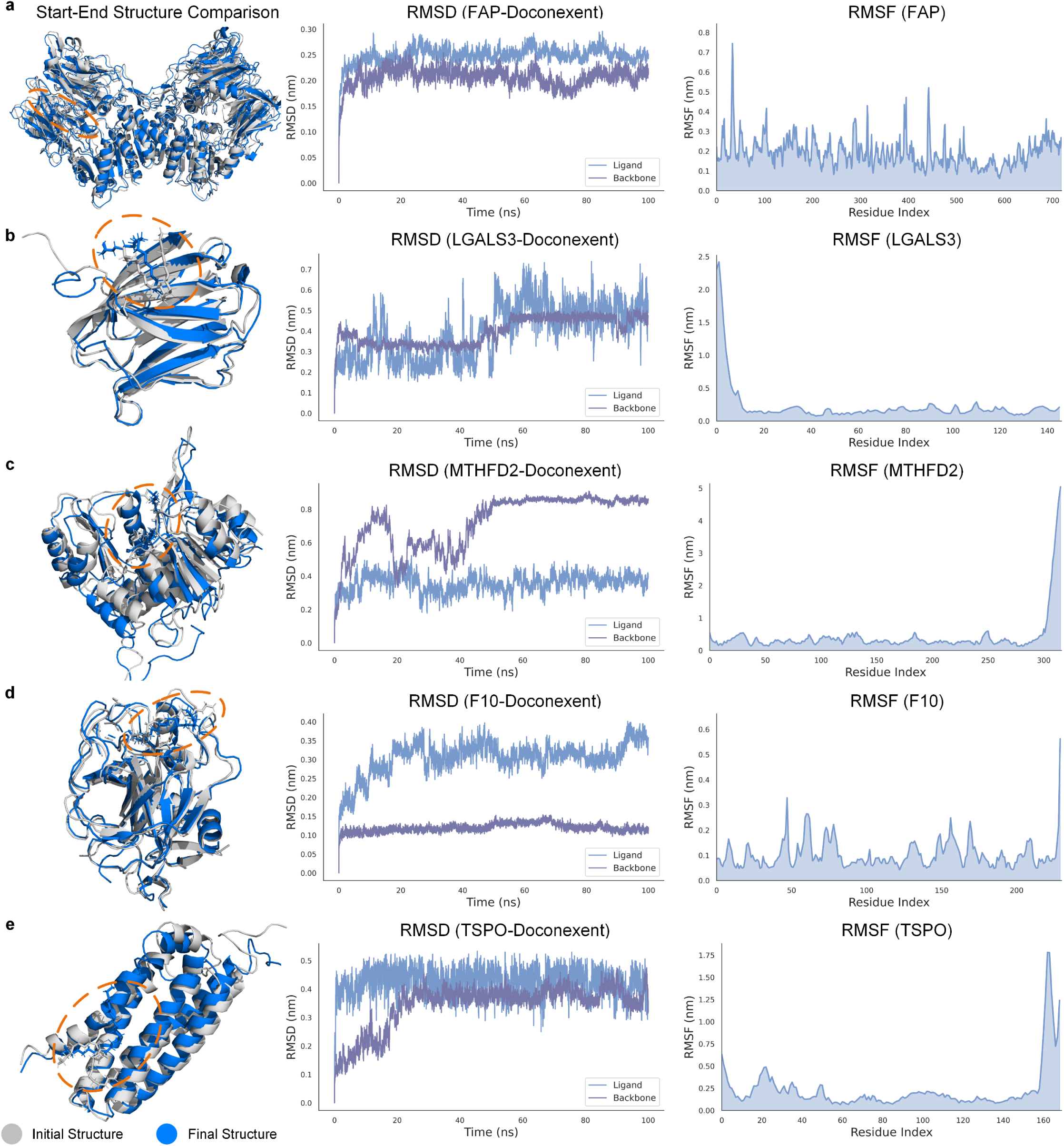
Molecular dynamics simulation analysis of Doconexent-target protein complexes. **a-e,** Molecular dynamics analysis of five protein-target complexes showing: (left) structural comparison between initial (gray) and final (blue) conformations with orange circles highlighting the binding pocket, (middle) RMSD trajectories over 100ns simulation for both ligand and protein backbone, and (right) RMSF analysis showing residue flexibility for (a) FAP complex, (b) LGALS3 complex, (c) MTHFD2 complex, (d) F10 complex, and (e) TSPO complex. All simulations were performed at 310.15K and 1 atm pressure. Crystal structures for FAP, LGALS3, MTHFD2, and F10 were obtained from RCSB Protein Data Bank, while the TSPO structure was generated using AF3.

Trajectory analysis through Root Mean Square Deviation (RMSD) and Root Mean Square Fluctuation (RMSF) calculations (Fig. 7, middle and right panels) provided detailed insights into the binding stability. As shown in Fig. 7a, the FAP-Doconexent complex showed rapid initial RMSD increases for both ligand and backbone, stabilizing around 0.2-0.25 nm throughout the simulation, with the ligand showing slightly higher fluctuations than the backbone, indicating achievement of a stable equilibrium state.

LGALS3 and MTHFD2 complexes with Doconexent (Fig. 7b-c) demonstrated distinct behaviors. The LGALS3 complex showed initial stability followed by increased ligand RMSD fluctuations (0.2-0.3 nm) after 40 ns, while its backbone maintained relatively stable RMSD around 0.2 nm. The MTHFD2 complex exhibited a gradual increase in ligand RMSD, reaching approximately 0.4 nm, with backbone RMSD stabilizing around 0.3 nm. Their RMSF analyses revealed localized flexibility, with LGALS3 showing distinct peaks in N-terminal residues (0-15, ∼2.5 nm) and MTHFD2 displaying significant C-terminal mobility (residues 300-320, ∼5 nm). These terminal region fluctuations likely reflect the absence of structural constraints in the crystal structures rather than instability in the protein-ligand binding interface.

The F10-Doconexent complex (Fig. 7d) showed gradual increases in ligand RMSD, reaching around 0.35 nm, while its backbone maintained relatively stable RMSD values around 0.15 nm. Despite these variations, the final binding pose closely resembled the initial configuration, confirming stable pocket retention. The RMSF profile showed moderate fluctuations distributed across various regions of the protein.

The TSPO-Doconexent complex (Fig. 7e) exhibited stable backbone RMSD around 0.2 nm, while the ligand RMSD showed larger fluctuations around 0.25-0.3 nm. The RMSF analysis revealed a notable peak around residues 150-170, which likely stems from lower confidence predictions in these regions of the AF3-generated structure, rather than inherent binding instability.

Collectively, our MD simulation analyses provide multiple supporting evidence for the stable binding of Doconexent to these five PDAC-associated targets. The maintenance of binding poses throughout the simulations and the convergence of RMSD values demonstrate stable protein-ligand interactions. Notably, the observed structural fluctuations were primarily localized to peripheral regions, while the core binding interfaces remained stable, as revealed by RMSF analyses. These computational findings provide a theoretical foundation supporting Doconexent’s further development as a potential PDAC therapeutic lead compound.

## Discussion

In this study, we introduced scKAN, a knowledge distillation framework that addresses key challenges in single-cell analysis through three major innovations: efficient knowledge transfer from single-cell LLMs, interpretable cell-type-specific gene set identification through activation curves, and cell-type-specific marker gene discovery via importance scores. Our comprehensive evaluation across four diverse datasets demonstrated scKAN’s superior performance in cell-type annotation, while its application to PDAC drug discovery validated its practical utility in translational research.

The technical advances of scKAN represent a significant step forward in computational efficiency for single-cell analysis. Unlike current LLMs, scKAN achieves an average 1.06% improvement in accuracy and 6.63% improvement in macro F1 score with reduced resource requirements through its lightweight KAN architecture. This efficiency gain is particularly valuable for widespread adoption in biomedical research settings where computational resources may be limited. Moreover, the interpretability provided by KAN’s learnable univariate functions offers unique insights into cell-gene relationships that are more biologically intuitive than Transformer-based attention mechanisms.

The biological insights derived from scKAN’s gene set identification capability have demonstrated both established and novel biological relationships. Our analysis identified functionally related gene clusters enriched in known biological pathways, particularly in immune cell development and pancreatic cell differentiation. These findings were further validated through pathway enrichment analysis, showing significant overlap with established molecular signatures. Notably, scKAN’s ability to capture subtle gene-gene relationships through activation curve similarities revealed previously unrecognized functional connections, suggesting potential new regulatory mechanisms in cell-type-specific contexts.

scKAN’s marker gene identification capability demonstrated remarkable accuracy across diverse cell types, combining both statistical robustness and biological relevance. The model successfully captured both strongly and subtly differentially expressed genes, outperforming SOTA methods when selecting limited candidate markers. The model’s importance scores strongly correlate with differential expression patterns while capturing more complex regulatory relationships, providing a more nuanced understanding of cell-type-specific gene expression programs. This capability was particularly valuable in identifying both canonical markers and novel candidates, as demonstrated by our validation of key markers such as SEL1L3 in naïve B cells and CST7 in NK cells.

The application of scKAN to PDAC research demonstrates its potential impact on therapeutic discovery. Our identification of ductal cell-specific markers led to the discovery of novel drug targets, with Doconexent emerging as a promising candidate. The stability of predicted drug-target interactions, validated through docking and molecular dynamics simulations, establishes a robust computational drug repurposing pipeline. This success highlights how scKAN’s interpretable framework can bridge the gap between single-cell analysis and therapeutic development, offering a new paradigm for targeted drug discovery.

From a methodological perspective, scKAN advances single-cell analysis by introducing a framework that balances computational efficiency with biological interpretability. This efficiency is demonstrated by a 15.8-fold reduction in GPU memory usage and 5.4-fold faster training compared to the teacher model scGPT, while maintaining robust performance. The integration of knowledge distillation with interpretable architecture provides a template for future developments in computational biology, potentially extending beyond single-cell analysis to other high-dimensional biological data types. While our current implementation utilizes scGPT as the teacher model, the framework’s design allows for seamless integration of more advanced LLMs.

Despite these promising results, several limitations and opportunities for future development warrant discussion. First, while scKAN demonstrates potential in cell-type annotation and marker gene identification, its naive KAN framework architecture currently limits its application in more complex single-cell analysis tasks such as perturbation response prediction and multi-modal data integration. Second, the comparable performance between KAN networks and MLPs on specific datasets suggests room for architectural refinement to enhance representational capacity while maintaining interpretability. Third, although scKAN effectively identifies potential therapeutic targets for specific diseases, the current drug repurposing workflow still relies on external methods for compound lead screening, indicating the need for an end-to-end solution from gene signatures to disease-specific drug candidates. Future work should address these limitations by extending the KAN framework with graph-based architectures for complex analysis tasks, incorporating multimodal learning capabilities for integrated omics analysis, and developing an end-to-end solution from target identification to lead compound screening.

Overall, the broader impact of scKAN extends beyond its immediate technical contributions. By providing an interpretable and efficient framework for single-cell analysis, it empowers researchers to extract meaningful biological insights from increasingly large and complex datasets. The success in PDAC drug discovery suggests potential applications in precision medicine across various diseases, including but not limited to cancer, autoimmune disorders, and neurodegenerative diseases, where cell-type-specific therapeutic targeting could improve treatment efficacy. Moreover, the framework’s ability to identify novel gene-gene relationships and drug targets could accelerate the drug discovery process, potentially reducing the time and cost of therapeutic development.

## Methods

### Kolmogorov–Arnold Networks

The architecture of KAN^21^ leverages the Kolmogorov-Arnold representation theorem, which states that any continuous multivariate function *f* can be represented as a composition of a finite number of continuous univariate functions:

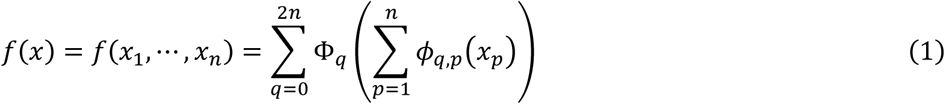

where *ϕ*_*q*,*p*_: [0,1] → ℝ, Φ_*q*_: ℝ → ℝ. Unlike traditional neural networks, such as MLPs that use fixed activation functions, KAN uses learnable activation functions on the network edges. In this design, each weight parameter in KAN can be replaced by a univariate function, which is typically parameterized using spline functions. This approach provides high flexibility and can model complex functions with fewer parameters, which improves model interpretability.

Specifically, KAN uses multiple layers of learnable activation functions as network edges. In contrast to MLPs, KAN does not apply nonlinear Transformations when aggregating function outputs:

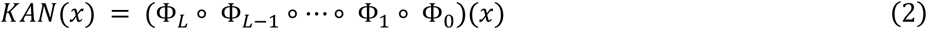

The activation functions are characterized through spline functions:

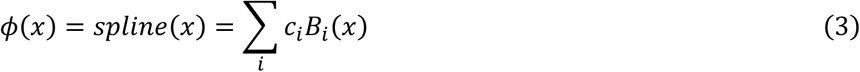

where *c*_*i*_ are learnable parameters and *B*_*i*_ (*x*) are B-spline basis functions defined on a grid with density controlled by a hyperparameter interval *G*. Using a larger *G* allows more control over the spline and higher precision but requires learning more parameters. The flexibility of splines enables the network to adaptively model complex relationships in data by adjusting their shapes. It minimizes approximation errors and enhances the network’s ability to learn subtle patterns from high-dimensional datasets.

In addition to spline functions, previous studies^84^ have shown that the Radial Basis Function (RBF) can also be used to approximate the activation function ϕ(*x*) in KAN:

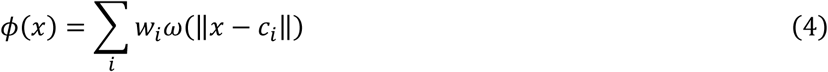

where *w*_*i*_ are learnable weights and ω is the radial basis function that depends on the distance between input *x* and center *c*_*i*_. The Gaussian function is the most common choice for RBFs:

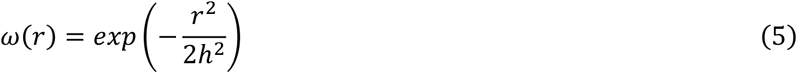

where *r* is the radial distance and ℎ is a hyperparameter that controls the spread of the function.

### Single-cell Large Language model

In this study, we use scGPT^18^, a SOTA foundation model for single-cell analysis, as the teacher model. We aim to transfer the representation patterns learned by scGPT, which was pre-trained on 33M single-cell data, into the lightweight scKAN model. scGPT consists of multiple self-attention Transformer layers that process tokenized genes and their expression values to extract features.

Specifically, for single-cell data containing N cells and M genes, the expression matrix can be represented as: *X* ∈ ℝ^*N*×*M*^, where each element *x_ij_* represents the RNA molecule read count for scRNA-seq data or chromatin accessibility of a peak region for scATAC-seq data. The model then obtains gene embeddings through gene tokens:

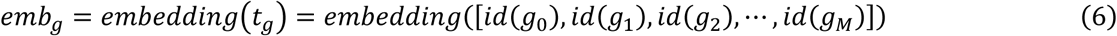

Additionally, the expression values are divided into B continuous intervals through binning: [*b*_*k*_, *b*_*k*+1_], where *k* ∈ {1,2, …, *B*}. The model then obtains expression embeddings through binning value tokens:

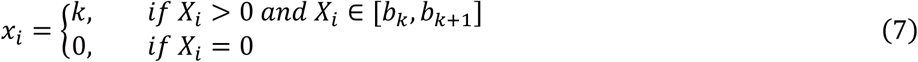

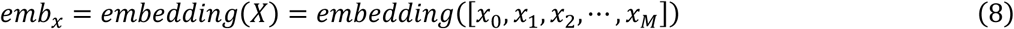

The model also incorporates conditional tokens containing various meta-information related to individual genes, such as cell batch (represented by batch tokens). We use input vectors with the same dimension as the input genes to represent position-aware conditional tokens and perform embedding. This embedding is represented as:

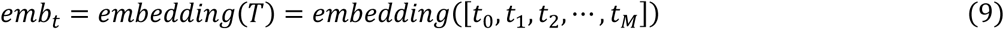

The embeddings are then concatenated and fed into Transformer layers to extract high-dimensional features:

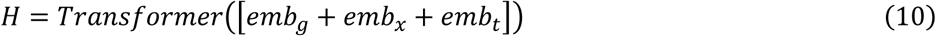

For cell-type annotation tasks, cell-level features can be obtained through the ‘[CLS]’ token, and class predictions are made using an MLP:

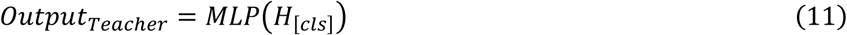

### Distillation Learning Framework

To enhance the efficiency of cell-type annotation, we propose a knowledge distillation framework with domain-aware learning principles that transfers the cell-type annotation capability from a pre-trained LLM (teacher model) to a lightweight model (student model, scKAN). We design a comprehensive loss function system comprising three components to ensure effective knowledge transfer and maintain annotation accuracy.

First, we implement a knowledge distillation loss to transfer the learned representations from the teacher model to the student model. This loss combines the conventional cross-entropy loss with a soft target distribution from the teacher model:

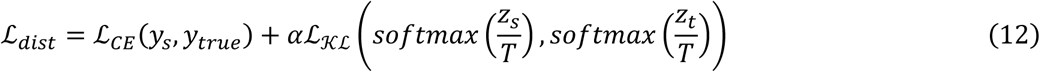

where *z*_*s*_ and *z*_t_ denote the logits from student and teacher models, respectively. T is the temperature parameter that controls the softness of probability distribution, and α balances the contribution of the distillation term.

Second, we introduce a self-entropy loss to maintain model robustness. In cell-type annotation, models may sometimes over-concentrate on certain dominant cell types while neglecting rare cell populations. It leads to a ‘trivial solution’ in which the model’s predictions are heavily biased towards specific cell types. To prevent this, we employ a self-entropy loss that encourages a balanced distribution of predictions across different cell types:

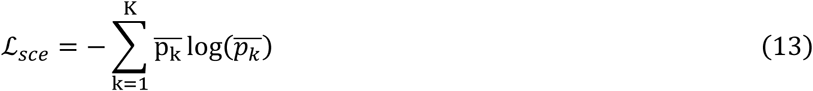

where 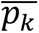 represents the mean probability of class *k* across the batch, and *K* is the number of cell types. This loss term ensures the model maintains sensitivity to all cell types, including rare populations, by penalizing highly skewed prediction distributions.

Third, we incorporate a DDC loss to enhance the quality of learned cell-type representations. The DDC loss was initially proposed with three components: feature-assignment consistency (ℒ_1_), assignment orthogonality (ℒ_2_), and feature-target consistency (ℒ_3_). In our adaptation, we modify the original DDC loss by removing the ℒ_2_ term, which enforces orthogonality between different cluster assignments through L2 regularization. We omit this term because, in our cell-type annotation task, the orthogonality constraint is inherently satisfied through the knowledge distillation process, where the teacher model already provides well-separated cell type representations. Therefore, our modified DDC loss focuses on two essential components:

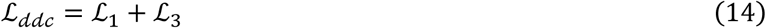

The first component, ℒ_1_, measures the consistency between hidden features and cluster assignments using Cauchy-Schwarz divergence:

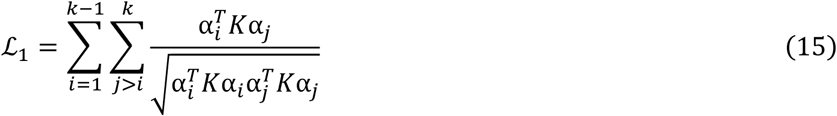

Here, *K* is the kernel matrix computed from hidden features using a Gaussian kernel:

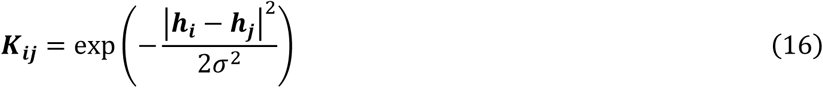

where ℎ_*i*_ represents the hidden features of sample *i*, and σ is the Gaussian kernel bandwidth determined based on the median distance in the batch. The second component, ℒ_3_, optimizes the relationship between cluster assignments and ideal one-hot distributions using a similar formulation with an exponential distance matrix.

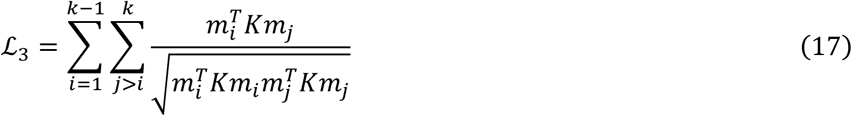

Where ***m_i_*** = [*m_q,i_*] ∈ *R^n^* and 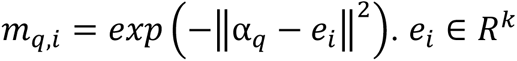 is a vector denoting the *i*th corner of the simplex, representing the *i*th cluster.

The complete training objective combines these three losses to simultaneously achieve knowledge transfer, maintain prediction balance, and optimize cell-type-specific feature representations:

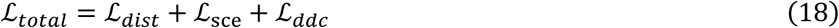

This comprehensive loss function design enables our lightweight model to effectively learn from the teacher model while maintaining robust and balanced cell-type annotation capabilities.

### Dataset

PBMC 10k Dataset^29^: The Peripheral Blood Mononuclear Cells (PBMC) 10K dataset was generated from a healthy donor using the 10x Genomics single-cell multiome ATAC + gene expression platform (Cell Ranger ARC 1.0.0). It comprises 9,631 cells, comprising 19 major immune cell populations, including CD4+ T cells, CD8+ T cells, and naïve B cells. The dataset is available at 10X Genomics official website (https://support.10xgenomics.com/single-cell-multiome-atac-gex/datasets/1.0.0/pbmc_granulocyte_sorted_10k). In this study, we focused on analyzing the gene expression (scRNA-seq) component of this multiome dataset.

Muto2021 Kidney Dataset^30^: This dataset, accessed from the Gene Expression Omnibus database under accession number GSE151302, represents a multi-omics study of adult human kidney cells, combining single-nucleus ATAC-seq (snATAC-seq) and RNA-seq (snRNA-seq) data. It provides detailed insights into kidney cell heterogeneity, focusing on proximal tubule epithelial cells. The dataset contains 19,985 cells with 13 distinct cell types identified. In this study, we focused on analyzing the RNA-seq component of this multi-omics dataset.

Human Pancreas Dataset^14^: This dataset integrates scRNA-seq data from five independent studies of human pancreas cells accessible from the Gene Expression Omnibus (GEO) database^85^. The data was strategically split into reference and query sets based on data sources. The reference set (10,600 cells) consists of two studies: Baron et al.^86^ (GSE84133) and Muraro et al.^87^ (GSE85241). The query set (4,218 cells) comprises three studies: Xin et al.^88^ (GSE81608), Segerstolpe et al.^89^ (E-MTAB-5061), and Lawlor et al.^90^ (GSE86473). The reference set encompasses 13 distinct cell populations, while the query set contains 11 cell types.

Myeloid Dataset^31^: The myeloid dataset was retrieved from the GEO database under accession number GSE154763, spanning nine cancer types. The reference set contains 9,748 cells from six cancer types: KIDNEY, LYM, PAAD, THCA, UCEC, and cDC2. The query set includes 3,430 cells from three types of cancer: ESCA, MYE, and OV-FTC. This dataset was preprocessed by Cui et al.^18^, resulting in an initial feature set of 3,000 genes. The dataset mainly focuses on myeloid cell populations across different cancer contexts.

Data Processing: Gene expression matrices were preprocessed using the Scanpy library^91^. The 2000 most highly variable genes were identified using the Seurat v3^92^. Raw counts were stored separately, followed by total-count normalization and log-transformation of the data. The normalized data underwent scaling to unit variance and zero mean. Principal component analysis (PCA) was performed for initial dimensionality reduction. A neighborhood graph was constructed using the cosine metric on the PCA representation, which was subsequently used for UMAP embedding generation.

### Baseline

Seurat^8^: A widely-used analysis pipeline that follows standard preprocessing steps, performs clustering analysis, and assigns cell type labels to clusters using established marker gene expressions.

CellTypist^10^: An annotation method integrating a reference database of immune cell types from multiple public datasets. The approach implements regularized linear models with stochastic gradient descent for automated cell-type identification. Through analysis of immune populations across different tissues, CellTypist captures both tissue-specific and shared features of immune cell subsets.

Cellcano^13^: A supervised learning framework designed for scATAC-seq data analysis. The method uses a two-stage learning strategy to reduce the impact of data distribution differences between reference and query datasets. The approach performs well across multiple cell typing tasks while maintaining computational efficiency.

TOSICA^14^: A Transformer-based model that incorporates biological knowledge into cell-type annotation. TOSICA provides accurate cell type labels and biological interpretations by analyzing pathway and regulon information. The method demonstrates effectiveness in identifying cell states during disease progression.

Geneformer^17^: A deep learning model that learns from an extensive collection of single-cell transcriptomes using attention mechanisms. The approach captures gene regulatory relationships through self-supervised training and allows adaptation to specific biological tasks with small amounts of data.

scGPT^18^: A large-scale generative model that processes cellular gene expression patterns using Transformer architecture. Like language processing systems, scGPT learns the relationships between genes across millions of cells. The model supports multiple downstream tasks through transfer learning, including cell annotation and data integration.

### Training Settings

The training consists of two main stages: LLM fine-tuning and knowledge distillation to scKAN.

We employed five-fold cross-validation for model evaluation. The data was initially split into reference and query sets with a ratio of 8:2. The reference set was further divided into training and validation sets at 7:1, resulting in a final split ratio of 7:1:2.

In the first stage, the LLM was fine-tuned for 10 epochs with a learning rate 5e-5. The number of bins for continuous value discretization was set to 51. The Adam optimizer was employed with an epsilon value of 1e-4. A StepLR scheduler was implemented to adjust the learning rate during training with predefined intervals and decay ratio.

For the knowledge distillation stage, scKAN was trained for 10 epochs with a higher learning rate of 5e-4. The knowledge distillation process used a temperature of 10 and combined soft and hard targets with weights of 0.5. All experiments were conducted on NVIDIA RTX 3090 GPUs.

The model performance was evaluated using multiple metrics: accuracy, macro F1-score, precision, and recall. These metrics were chosen to comprehensively assess overall annotation accuracy and performance across individual cell types. All reported results represent the mean and standard deviation across the five-fold cross-validation. Detailed descriptions of the training settings and hyperparameter configurations can be found in Supplementary Note 1.

### Gene Set Identification Using scKAN

To identify cell-type-specific gene sets, we developed a systematic approach leveraging the trained scKAN model’s interpretable architecture. We employed a two-layer scKAN structure to establish a direct mapping between cell types and gene expression patterns, enabling the extraction of cell-gene activation curves. This process consists of three main steps: curve extraction, gene set optimization, and enrichment analysis.

First, we extracted the learned curves from the final layer of scKAN for each cell type using the ‘plot_curve’ method. These curves represent the learned relationships between gene expression and cell type. Specifically, for each gene *i* and cell type *j*, Specifically, for each gene *i* and cell type *j*, we obtained a curve by sampling 400 points within the range ‘[-2-2h, 2+2h]’, where ℎ is the grid interval of the radial basis functions.

We then employed a beam search algorithm to identify optimal gene sets for each cell type. The objective function was defined as the sum of pairwise cosine similarities between the curves of genes within a set:

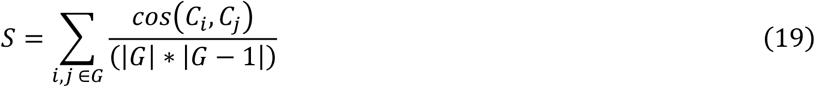

where *G* represents a candidate gene set, *Ci* and *Cj* are the curves for genes *i* and *j*, and |*G*| is the size of the gene set. The beam search maintained the top-k partial solutions at each step, where *k* was set to 20, and the final gene set size was limited to 20 genes per cell type.

The identified gene sets were subsequently analyzed using GSEA implemented through the GSEApy package^93^. We specifically focused on immune-related pathways using the immune gene set from molecular signatures database^42,94^. This analysis provided insights into the immunological processes and functions associated with cell-type-specific gene expression patterns.

### Identification of Cell-type-specific Marker Genes

To identify markers that characterize different cell types, we utilized the attribution mechanism initially designed for network pruning in KAN. The computation is implemented using the original KAN architecture to preserve the network’s attribution capabilities. The attribute function generates edge scores through backward propagation from the output layer to the input layer. The scores are initialized to an identity matrix representing cell type outputs. Then, the function computes successive transformations through node-to-subnode and subnode-to-edge connections, producing attribution scores reflecting gene-cell type associations. Detailed implementation of the attribution mechanism can be found in the original paper^21^.

These attribution scores are represented as a matrix of dimensions N × M, where N represents the number of cell types and M represents the number of genes. Each element (i,j) in this matrix quantifies the contribution strength of gene j to the cell type i. The resulting edge scores provide a quantitative measure of gene importance for each cell type, enabling the identification of cell-type-specific marker genes. Higher attribution scores indicate stronger associations between genes and specific cell types, revealing potential marker genes for each cell population^21^.

### ADMET Property Evaluation

To evaluate the drug-likeness and potential pharmacological properties of our screened lead compounds, we performed comprehensive Absorption, Distribution, Metabolism, Excretion, and Toxicity (ADMET) analyses using ADMETlab 3.0^76^, a widely used web platform for systematic ADMET property prediction.

Multiple physicochemical properties were assessed for each compound. These include molecular weight (MW), which influences drug distribution and elimination; partition coefficient (LogP) and distribution coefficient (LogD), which reflect lipophilicity and drug partitioning within the body; and aqueous solubility (LogS), which is crucial for oral bioavailability. We also examined structural features, including the number of hydrogen bond acceptors (nHA) and donors (nHD), which affect solubility and receptor binding; topological polar surface area (TPSA), which correlates with drug transport properties; and the number of rotatable bonds (nRot), which influences oral bioavailability. Additional structural characteristics such as the number of rings (nRing), maximum ring size (MaxRing), number of heteroatoms (nHet), formal charge (fChar), and number of rigid bonds (nRig) were analyzed to evaluate drug-receptor interactions and overall pharmacokinetics.

These comprehensive predictions provide valuable insights into the compounds’ potential as drug candidates and help identify possible limitations in their pharmacological profiles.

### Molecular Docking Analysis

To investigate potential interactions between screened compounds and target proteins, we performed molecular docking simulations using CB-Dock2^77^. This advanced protein-ligand blind docking tool integrates cavity detection, docking, and homologous template fitting. We set the maximum number of binding cavities for each protein-ligand pair to 5 to ensure comprehensive coverage of potential binding sites while maintaining computational efficiency.

CB-Dock2 first detects potential binding cavities on the protein surface using a curvature-based approach. It then performs molecular docking by combining structure-based and template-based docking strategies. The structure-based docking employs AutoDock Vina to sample ligand conformations within the detected cavities, while the template-based approach leverages known protein-ligand complex structures to guide the docking process. Multiple binding poses were generated and ranked for each cavity based on their binding scores. The docking results provide insights into the potential binding modes and interaction patterns between the screened compounds and their target proteins. The top-ranked protein-ligand complexes were subsequently used as initial configurations for molecular dynamics simulations to evaluate the stability of these interactions.

### Molecular Dynamics Simulation

We employed molecular dynamics (MD) simulations to investigate protein-ligand complex dynamics. The system was prepared using the OpenFF Toolkit and OpenMM framework^95^. The protein was parameterized using the AMBER ff14SB force field, while the ligand parameters were derived using the GAFF-2.11 force field. The complex was solvated in an explicit TIP3P water box extending 10 Å from the solute surface. Counter-ions (Na+ and Cl-) were added to neutralize the system and achieve physiological ionic strength.

The system underwent energy minimization followed by a two-stage equilibration protocol: a 50,000-step NVT equilibration at 310.15 K using a Langevin thermostat (collision frequency of 1.0 ps⁻¹), followed by a 50,000-step NPT equilibration at 1 atm pressure using a Monte Carlo barostat. Production MD simulations were performed with a 2 fs time step, with all bonds involving hydrogen atoms constrained using the SHAKE algorithm. The temperature was maintained at 310.15 K using a Langevin thermostat. The production run was extended to 100 ns, and coordinates were saved every 1000 steps for analysis.

All simulations were performed using OpenMM^95^ on GPU-accelerated computing resources with periodic boundary conditions and Particle Mesh Ewald for long-range electrostatic interactions.

Trajectory analysis was performed using the MDTraj^96^ library to evaluate system stability and conformational changes. We calculated the RMSD of the ligands and binding site residues and the RMSF of the protein backbones. The RMSD of molecular coordinates indicates conformational changes over time and is calculated as:

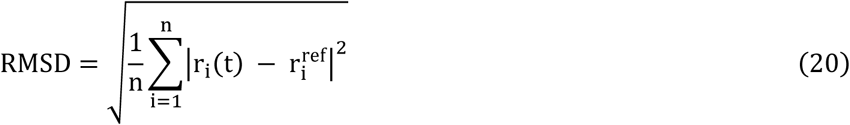

where n is the number of atoms, *r_i_*(*t*) is the position of atom *i* at time *t*, and *r_i_*^*ref*^ is the position of atom *i* in the initial complex structure obtained from molecular docking.

The RMSF measures the fluctuation of each residue around its average position during the simulation, calculated as:

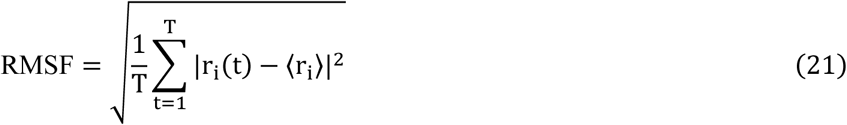

where T is the total number of time steps in the simulation, *r_i_*(*t*) is the position of residue *i* at time *t*, and ⟨*r_i_*⟩ is the average position of residue *i* over the entire simulation.

## Data Availability

All datasets used are obtained from public data repositories. The myeloid dataset is publicly accessible from the GEO database using accession number GSE154763. The processed human pancreas dataset was retrieved from https://github.com/JackieHanLab/TOSICA. The PBMC 10k dataset is available at https://support.10xgenomics.com/single-cell-multiome-atac-gex/datasets/1.0.0/pbmc_granulocyte_sorted_10k. The Muto-2021 dataset is available at GEO with the accession number GSE151302.

## Code Availability

The codebase for this study is publicly available at: https://github.com/hehh77/scKAN.

## Author Contributions Statement

H.H.: Methodology, Conceptualization, Software, Validation, Writing - original draft, Visualization. Z.T.: Conceptualization, Formal analysis, Investigation. G.C.: Visualization, Investigation. F.X.: Data Curation, Formal analysis. Y.H.: Formal analysis, Data Curation. Y.F.: Validation, Formal analysis. J.W.: Supervision, Funding acquisition, Resources. Y.H: Writing - Review & Editing, Funding acquisition. Z.H.: Writing - Review & Editing, Funding acquisition, Resources, Supervision. K.T.: Writing - Review & Editing, Supervision, Funding acquisition.

## Competing interests Statement

The authors declare no competing interests.

## Notes

### Competing Interest Statement

The authors have declared no competing interest.

